# Disentangling neocortical alpha/beta and hippocampal theta/gamma oscillations in human episodic memory formation

**DOI:** 10.1101/2020.01.22.915330

**Authors:** Benjamin J. Griffiths, María Carmen Martín-Buro, Bernhard P. Staresina, Simon Hanslmayr

## Abstract

To form an episodic memory, we must first process a vast amount of sensory information about a to-be-encoded event and then bind these sensory representations together to form a coherent memory. While these two cognitive capabilities are thought to have two distinct neural origins, with neocortical alpha/beta oscillations supporting information representation and hippocampal theta-gamma phase-amplitude coupling supporting mnemonic binding, evidence for a dissociation between these two neural markers is conspicuously absent. To address this, seventeen human participants completed a sequence-learning task that first involved processing information about three stimuli, and then binding these stimuli together into a coherent memory trace, all the while undergoing MEG recordings. We found that decreases in neocortical alpha/beta power during sequence perception, but not mnemonic binding, correlated with enhanced memory performance. Hippocampal theta/gamma phase-amplitude coupling, however, showed the opposite pattern; increases during mnemonic binding (but not sequence perception) correlated with enhanced memory performance. These results demonstrate that memory-related decreases in neocortical alpha/beta power and memory-related increases in hippocampal theta/gamma phase-amplitude coupling arise at distinct stages of the memory formation process. We speculate that this temporal dissociation reflects a functional dissociation in which neocortical alpha/beta oscillations could support the processing of incoming information relevant to the memory, while hippocampal theta-gamma phase-amplitude coupling could support the binding of this information into a coherent memory trace.

## Introduction

An episodic memory is a personal detail-rich, long-term memory that is anchored to a unique point in time and space (Tulving, 2002). The formation of these memories are thought to rely on both neocortical alpha/beta and hippocampal theta/gamma oscillations (Hanslmayr et al., 2016), both of which are prevalent in a wide range of human episodic memory tasks (for reviews, see Hanslmayr & Staudigl, 2014; Nyhus & Curran, 2010).

Neocortical alpha/beta desynchrony is thought to be beneficial for information representation (Hanslmayr et al., 2012). This idea is derived from the tenets of information theory, which propose that unpredictable states (e.g., a desynchronised network, where the firing of one neuron cannot predict the firing of another) convey substantially more information than predictable states. In direct support of this idea, neocortical alpha/beta (8-20Hz) power decreases have been shown to correlate with the enhanced fidelity of neural representations present in BOLD signal (Griffiths, Mayhew, et al., 2019). Moreover, interfering with these power decreases via transcranial magnetic brain stimulation impairs episodic memory formation (Hanslmayr et al., 2014). Together, these findings (see also Fellner et al., 2013; Griffiths et al., 2021; Karlsson et al., 2020; Long & Kahana, 2015; Martín-Buro et al., 2020; Sederberg et al., 2007) suggest that alpha/beta power decreases are intimately tied to the successful representation of information pertaining to episodic memories.

Hippocampal theta and gamma oscillations also play a pivotal role in episodic memory formation (e.g. Bahramisharif, Jensen, Jacobs, & Lisman, 2018; Heusser, Poeppel, Ezzyat, & Davachi, 2016; Staudigl & Hanslmayr, 2013; Tort, Komorowski, Manns, Kopell, & Eichenbaum, 2009). The phase of theta is thought to determine whether long-term potentiation (LTP) or long-term depression (LTD) occurs (Hasselmo et al., 2002), and gamma synchronisation compliments this process by driving neurons to fire at the frequency optimal for spike-timing dependent plasticity (STDP; Bi & Poo, 1998; Jutras, Fries, & Buffalo, 2009; Nyhus & Curran, 2010). By combining these two phenomena, hippocampal theta-gamma phase-amplitude coupling is well-suited for mnemonically binding disparate sources of information into a coherent memory trace (Hanslmayr et al., 2016; Lisman & Jensen, 2013).

On a cognitive level however, many paradigms probing human episodic memory formation involve substantial overlap in information representation and mnemonic binding, making it difficult to conclude that their associated neural phenomena are truly dissociable. Here, we addressed this problem by using a paradigm that invokes a temporal shift in the ratio of these cognitive processes. Seventeen participants were briefly presented with a sequence of three stimuli (always consisting of an object, a feature and a scene), and then given a small window to intentionally bind these stimuli together for a later associative memory test^*^. We hypothesised that memory-related changes in neocortical alpha/beta activity would show a distinct temporal dynamic to memory-related changes in hippocampal theta/gamma activity. Specifically, that (1) memory-related neocortical alpha/beta power decreases would be most prevalent during the perception of the sequence (from here on termed “sequence perception”), as this requires extensive processing of the details of each item prior to binding, and (2) memory-related increases in hippocampal theta-gamma phase-amplitude coupling would be most prevalent when participants intentionally associate the stimuli together (from here on termed “mnemonic binding”), given that theta-gamma coupling is a proxy for forms of long-term potentiation. Indeed, the results reported below support these hypotheses, suggesting that neocortical alpha/beta desynchrony and hippocampal theta/gamma synchrony arise at distinct stages of the memory formation process.

## Materials and Methods

### Participants

Twenty-eight participants were recruited (mean age = 25.4; age range = 20-33; 68% female; 82% right-handed). These participants received course credit or financial reimbursement in return for their participation. One participant was excluded for excessive head movement (greater than 2 standard deviations above group mean). Four participants were excluded for poor data quality (more than 50% of trials rejected for artifacts). Six participants were excluded for extreme memory performance (fewer than 15 trials in one of the three memory conditions). This left seventeen participants for further analysis (mean age = 24.9; age range = 20-32; 65% female; 82% right-handed). Ethical approval was granted by the Research Ethics Committee at the University of Birmingham (ERN_15-0335), complying with the Declaration of Helsinki.

### Experimental design

Each participant completed a visual associative memory task (see figure 1a). During encoding, participants were presented with a line drawing of an object, a pattern, and a scene (each for 1500ms, with a jittered 600ms (±100ms) fixation cross shown between each stimulus). Participants were then given a short interval to create a mental image incorporating the three stimuli to help them recall the stimuli for a later memory test. Participants were then asked to rate how difficult they found associating the triad. This question was used to keep participants attending to the task, rather than provide a meaningful metric for analysis. The next trial began after the participant had responded to the difficulty question. After associating 48 triads, participants started the distractor task. In the distractor task, participants attended to a fixation cross in the centre of a black screen. The fixation cross would flash momentarily (∼100ms) from light grey to either white or dark grey approximately every 20 seconds. The participants were instructed to count the number of times the fixation cross changed to white (ignoring the times it turned dark grey) and report this value at the end of the task (approximately 2.5mins later). The retrieval task followed the distractor. Here, participants were presented with the line drawing (for 3000ms) and asked to recall the mental image they made during the encoding phase. Then, participants were presented with three patterns (one correct and two lures) and asked to identify the pattern associated with the line drawing. After responding, participants were presented with three scenes (one correct and two lures) and again asked to identify the pattern associated with the line drawing. After responding, participants were then asked to indicate how confident they were about their choices. They could select ‘guess’ (i.e., they guessed their choice), ‘unsure’ (i.e. they could not remember the item, but had a feeling it was the correct choice), or ‘certain’ (i.e. they could vividly remember the item). Participants were asked to recall all 48 triads learnt in the earlier encoding phase.

**Figure 1.**
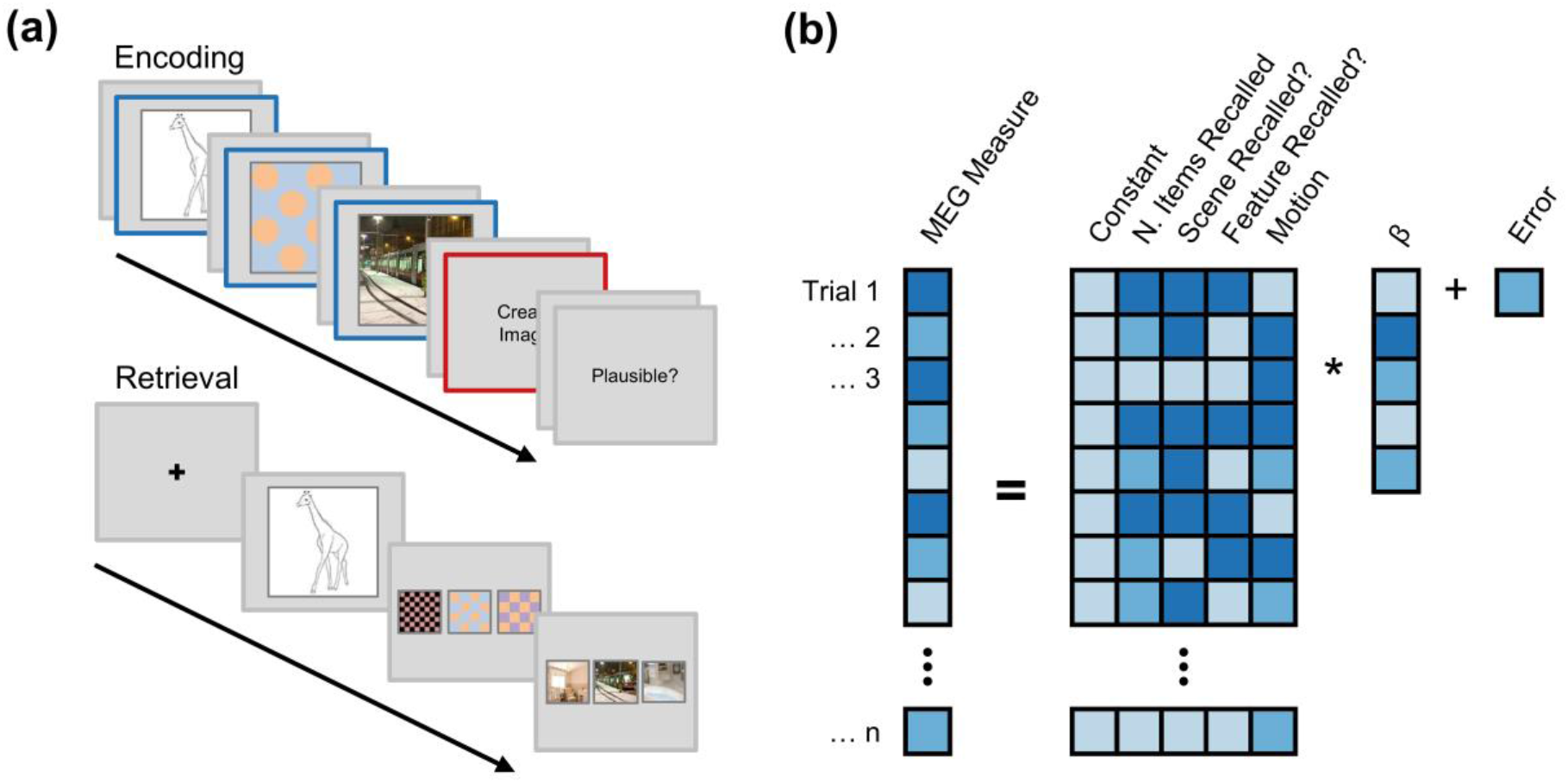
Overview of task and analytical approach. **(a)** Paradigm schematic. Participants were presented with a sequence of three visual stimuli. The sequence always began with a line drawing of an object, and was then followed by a pattern and a scene (each with a brief fixation cross shown between). Participants were then given a short interval to create a mental image incorporating the three stimuli, before being asked to rate how difficult it was to create the association. After a distractor task, participants were presented with the object as a cue and asked to recall both the pattern and the scene, each from a choice of three stimuli. After selection, participants had to rate how confident they felt about their response. The epochs representing “sequence perception” are outlined in blue, and the epoch representing “mnemonic binding” is outlined in red. **(b)** Analysis schematic. For each participant, spectral power and theta-gamma phase-amplitude coupling were modelled using a general linear model including predictors for the number of items recalled, scene and feature memory recall, and head motion. The resulting standardised beta coefficient for the central predictor (i.e., number of items recalled) was extracted, pooled across participants, and subjected to a one sample t-test to determine whether the number of items recalled predicted changes in spectral power and/or theta-gamma coupling.

Participants completed four blocks of this task (192 trials in total). The order in which the pattern and scene were presented during perception was swapped between each block (where a “block” is defined as a complete cycle of encoding, distractor and retrieval tasks). On blocks where scenes preceded patterns during perception, the presentation order at retrieval was also reversed.

For all responses, participants used two non-magnetic, single-finger optical response pads. The left pad allowed participants to cycle through the possible responses, and the right pad allowed participants to confirm their selection.

### Behavioural analysis

For each trial, memory performance was coded as either ‘complete’ (i.e., they remembered both the scene and the pattern), ‘partial’ (i.e. they remembered only one of the associates), or ‘forgotten’ (i.e. they remembered neither the scene nor the pattern). Any selection where the participant indicated that they guessed was marked as a ‘miss’.

### MEG acquisition

MEG data was recorded using a 306-channel (204 gradiometers, 102 magnetometers) whole brain Elekta Neuromag TRIUX system (Elekta, Stockholm, Sweden) in a magnetically shielded room. Participants were placed in the supine position for the duration of the experiment. Data was continuously recorded at a sampling rate of 1000Hz. The head shape of each participant (including nasion and left/right ear canal) was digitised prior to commencing the experiment. Continuous head position indicators (cHPI) were recorded throughout. The frequencies emitted by the cHPI coils were 293Hz, 307Hz, 314Hz and 321Hz. Magnetometer data was excluded from the main analysis as they contained substantial noise that could not be effectively removed or attenuated.

### MEG preprocessing

All data analysis was conducted in Matlab using Fieldtrip (Oostenveld et al., 2011) in conjunction with custom scripts. First, the data was lowpass filtered at 165Hz to remove the signal generated by the HPI coils. Second, the data was epoched around each event of interest. At encoding, the epochs reflected the time windows where each stimulus was presented (from here on termed ‘sequence perception’) and when the ‘associate’ prompt was presented (termed ‘mnemonic binding’). Sequence perception epochs began 2000ms before stimulus onset and ended 3500ms after onset (that is, 2000ms after stimulus offset). Mnemonic binding epochs began 2000ms before stimulus onset and ended 5000ms after onset (that is, 1500ms after stimulus offset). Third, independent components analysis was conducted, and any identifiable eye-blink or cardiac components were removed. Fourth, the data was visually inspected, and any artefactual epochs or sensors were removed from the dataset (mean percentage of trials removed: 18.0%; range: 5.7-32.2%).

### Movement correction

To identify participants with extreme head motion during MEG recordings, the recorded data was first highpass filtered to 250Hz to isolate the cHPI signal. Second, the variance of the signal for each sensor was computed across every time point of the continuous recording. Third, the variance was mean averaged across sensors to provide a singular estimate of change in cHPI signal across the duration of the experiment. Fourth, the mean variance and its standard deviation was calculated across participants. Lastly, participants with extreme head motion were identified as those with variance greater than two standard deviations above the group mean. These participants were excluded from further analysis.

To help attenuate motion-related confounds in the spectral power and phase-amplitude coupling analyses, a trial-by-trial estimate of motion was calculated. First, the data was highpass filtered at 250Hz. Second, the data was epoched into trials matching those outlined in the section above. Third, the envelope of the signal in each epoch was calculated (to avoid issues of mean phase angle difference in cHPI signal across trials). Fourth, the envelope was averaged over time to provide a single value for each epoch and channel. Fifth, the dot product was computed across sensors between the first epoch and every other epoch (algebraically: 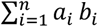, where *n* is the number of channels, ***a***_***i***_ is the power at sensor ***i*** during the first trial, and ***b***_***i***_ is the power at sensor ***i*** during the trial of interest). This provided a single value (between zero and infinity) for each trial that described how similar the topography of that trial was to the first trial – the higher the value, the more similar the topographies are between the two trials (with the assumption that the more dissimilar a cHPI topography is to the starting topography, the more the head has deviated from its starting position). These values were entered as a regressor of no interest in the central multiple regression analyses.

### Time-frequency decomposition and statistical analysis

Sensor-level time-frequency decomposition was conducted on the sequence perception and mnemonic binding epochs. For low frequencies, the preprocessed data was first convolved with a 6-cycle wavelet (−0.5 to 3 seconds [to 2 seconds for perceptual epochs to avoid the subsequent stimulus], in steps of 50ms; 2 to 40Hz; in steps of 1Hz). For high frequencies, Slepian multitapers were first used to estimate power (−0.5 to 3 seconds [to 2 seconds for perceptual epochs], in steps of 50ms; 40 to 100Hz, in steps of 4Hz). For this latter analysis, frequency smoothing was set to one quarter of the frequency of interest and temporal smoothing was set to 200ms. Second, planar gradiometers were combined by summing the power of the vertical and horizontal components. Third, for perceptual trials only, power was then averaged over the three stimulus presentation windows of each triad to provide mean power during perception of the triad. Any triads where one or more epochs had been rejected during preprocessing were excluded at this stage. We averaged spectral power across the three windows as we reasoned that this approach would be most sensitive to changes in spectral power that predicted the number of items later recalled. To successfully recall a stimulus, an alpha/beta power decrease must arise in two of the windows – the initial line drawing (i.e. the retrieval cue) and the to-be-recalled stimulus. As such, focusing analyses on a single stimulus is less sensitive to later memory performance than an aggregate measure created by averaging across the epochs. Fourth, the background 1/f characteristic was subtracted using an iterative linear fitting procedure.

To isolate oscillatory contributions, 1/f activity was attenuated in the time-frequency data by subtracting the linear fit of the 1/f characteristic (Griffiths, Parish, et al., 2019; Manning et al., 2009; Zhang & Jacobs, 2015). To this end, a vector containing values of each derived frequency (*A*) and another vector containing the power spectrum, averaged over all time-points and trials of the relevant memory condition (*B*) were log-transformed to approximate a linear function. The linear equation ***Ax = B*** was solved using least-squares regression, where ***x*** is an unknown constant describing the 1/f characteristic. The 1/f fit (***Ax***) was then subtracted from the log-transformed power spectrum (***B***). As this fit can be biased by outlying peaks (Haller et al., 2018), an iterative algorithm was used that removed probable peaks and then refitted the 1/f. Outlying peaks in this 1/f-subtracted power spectrum were identified using a threshold determined by the mean value of all frequencies that sat below the linear fit. The MEG power spectrum is the summation of the 1/f characteristic and oscillatory activity (i.e., at no point does oscillatory activity subtract from the 1/f), therefore all values that sit below the linear fit can be seen an estimate error of the fit. Any peaks that exceed the threshold were removed from the general linear model, and the fitting was repeated. Notably, as power for the low frequencies (2-40Hz) and high frequencies (40-100Hz) was calculated using different methods (wavelets and Slepian multitapers, respectively), the two bands have disparate levels of temporal and spectral smoothing. To avoid a spurious fitting due of the 1/f because of these differences, the 1/f correction was conducted separately for these two bands.

For statistical analysis, a trial-based multiple regression was run for each participant. Four regressors were used to predict observed power for every channel x frequency x time point independently. These four regressors were (1) number of items recalled, (2) whether the scene was recalled, (3) whether the pattern was recalled, (4) the change in head position [based on the motion calculation outlined above]. The first regressor was of primary interest, the second and third regressors isolated spectral power changes that can be uniquely described by whether the scene/pattern was recalled (respectively), and the fourth regressor accounted for changes in spectral power driven by head movement (see the next paragraph for notes on multicollinearity). The beta weight of the first regressor, obtained for a given channel x frequency x time point, was then standardised by dividing the standard error of the fit (providing a t-value) to attenuate the impact of poor model fits on the final analysis. Here, a positive standardised beta coefficient would indicate that spectral power increases with more items recalled, and a negative beta coefficient would indicate that spectral power decreases with more items recalled. The beta coefficients for each participant were pooled across the sample and entered into a one-tailed cluster-based permutation test (2000 permutations, alpha threshold = 0.05, cluster alpha threshold = 0.05, minimum neighbourhood size = 3; Maris & Oostenveld, 2007) to examine whether the observed fits consistently deviated from the null hypothesis (t=0) across participants. Clusters that produced a p-value less than 0.05 were considered significant. Cohen’s dz was used as the measure of effect size for these clusters (Lakens, 2013), where 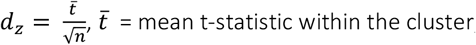, ***n*** = number of participants.

Notably, it is plausible to suggest that the three memory regressors are, to some extent, correlated and that this would introduce multicollinearity into the regression models. To test this, we calculated the Variance Inflation Factor (VIF) – a measure of the magnitude of multicollinearity. Rule of thumb suggests that a VIF greater than 10 is considered high and could compromise the model (Kutner et al., 2004). The VIF between number of items recalled and scene recall success was, on average, 2.615 (s.d. 2.621), and the VIF between number of items recalled and pattern recall success was, on average, 1.027 (s.d. 0.033). As these values fall below the threshold of 10, multicollinearity is not an apparent concern.

An additional analysis was conducted to confirm that alpha/beta power did indeed decrease following the onset of the sequence stimuli. This analysis followed the same approach as above (i.e., regression-based analyses across trials for each participant individually, and then group-level statistics on the resulting standardised beta coefficients). However, two changes were made: (1) the outcome variable became the change in 1/f-corrected spectral power from baseline (−250ms to stimulus onset; as opposed to raw spectral power), and (2) the memory-related regressors removed from the regression models, as they were of no relevance here.

### Source analysis

The preprocessed data was reconstructed in source space using individual head models and structural (T1-weighted) MRI scans for all but two individuals who did not wish to return for an MRI scan. For these two individuals, a standard head model and MRI scan was used (taken from the Fieldtrip toolbox; for details, see http://www.fieldtriptoolbox.org/template/headmodel). The head shape (together with the HPI coil positions) of each participant was digitised using a Polhemus Fasttrack system. The timelocked MEG data was reconstructed using a single-shell forward model and a Linearly Constrained Minimum Variance beamformer (LCMV; van Veen, van Drongelen, Yuchtman, & Suzuki, 1997). The lambda regularisation parameter was set to 1%.

### MEG phase-amplitude coupling computation and statistical analysis

For the phase-amplitude coupling analyses, we focused our analysis directly on source-reconstructed hippocampal virtual sensors. Given the depth and size of the hippocampus (it makes up around ∼1% of the MEG sourcemodel), the likelihood that legitimate hippocampal phase-amplitude coupling can be observed on the scalp is practically nil. Therefore, it makes most sense to move directly to source space and analyse source-localised measures of hippocampal activity.

To calculate the extent to which hippocampal gamma activity coupled to hippocampal theta phase, the modulation index (MI) was calculated (Tort et al., 2010). First, the peak theta and gamma frequencies were calculated by estimating power across all hippocampal virtual sensors (bilaterally, as defined by the automated anatomical labelling [AAL] atlas) using the same time-frequency decomposition method reported above^†^. The Matlab function *findpeaks()* was then used to extract the most prominent peak within the theta (2-7Hz) and gamma (40-80Hz) bands for each participant. Across participants, the mean theta peak was at 5.1Hz (standard deviation: 1.0Hz; range: 3.1-7.0Hz), and the mean gamma peak was at 66.1Hz (standard deviation: 4.6Hz; range: 59.0-73.0Hz) [see supplementary figure 1 for all plots]. Second, the time-series of the hippocampal virtual sensors were duplicated, with the first being filtered around the theta peak (±0.5Hz) and the second being filtered around the gamma peak (±5Hz). Third, the Hilbert transform was applied to the theta- and gamma-filtered time-series, with the phase of the former and power of the latter being extracted. Fourth, the time-series data was re-epoched, beginning 500ms after the onset of the stimulus/fixation cross and at the onset of the next screen. This attenuated the possibility that an event-related potential and/or edge artifacts from the filtering/Hilbert transform could influence the phase-amplitude coupling measure (Aru et al., 2014). Fifth, gamma power was binned into 12 equidistant bins of 30°, according to the concurrent theta phase. This binning was conducted for each trial and sensor separately. Sixth, the MI was computed by comparing the observed distribution to a uniform distribution. Seventh, the resulting MI values were subjected to a trial-based multiple regression conducted in the same manner as for the spectral power analyses. However, two additional regressors were added to this model: (1) hippocampal peak theta power [per trial, averaged across 500ms to 3000ms], (2) hippocampal peak gamma power [per trial, averaged across 500ms to 3000ms]. These regressors addressed the potential confound of concurrent power influence phase estimates (Aru et al., 2014). Eighth, these results were averaged over hippocampal virtual sensors and these per-participant standardised beta coefficients were subjected to a permutation-based one-sample t-test contrasting memory-related changes in phase-amplitude coupling to the null hypothesis (t=0). Notably, as we had focused our analyses on the peak theta and gamma frequencies, and used the average PAC values across virtual sensors, only a single statistical comparison was made. Therefore, no cluster-based multiple comparison correction was required. As the p-values reported here are estimated by permutation rather than from the parametric test, the t-values and degrees of freedom we report should only be used for reference.

We examined the spatial specificity of this effect by using the same pipeline as above to assess theta-gamma phase-amplitude coupling in the frontal, occipital, parietal, and temporal lobes (individually; as defined by the *wfupickatlas* toolbox for SPM). This analysis (plus source visualisation) help confirm that theta-gamma coupling observed in the hippocampus ROI originated from the hippocampus itself, rather than “bled in” from another region.

To compliment this analysis, we also ran a searchlight-based analysis. Here, we aimed to contrast the magnitude of the memory-related hippocampal coupling effect with memory-related coupling effects outside the hippocampus (in searchlights including approximately the same number of voxels; assuaging concerns that the lobe-based ROIs were too broad to detect local coupling effects). To this end, we iterated through every source voxel, identified its immediate neighbours (those immediately in front of and behind the voxel in 3-dimensional space [mini-cluster size: 27 voxels; for comparison, hippocampal ROI = 25 voxels]), and took the mean memory-related hippocampal phase-amplitude coupling within this mini-cluster. For each mini-cluster, this mean value was then contrasted against chance, and the resulting t-statistic was added to a distribution describing the magnitude of memory-related phase-amplitude coupling across the brain. A p-value was then dervied by comparing hippocampal coupling to the whole-brain distribution (as done in a permutation test), allowing us to infer the extent to which hippocampal phase-ampltiude coupling deviated from what was typical within the brain.

## Results

### Behavioural results

Participants, on average, correctly recalled both the associated pattern and associated scene on 38.3% of trials, recalled only one associated stimulus on 34.4% of trials, and failed to recall either associate on 27.3% of trials. Participants correctly recalled the associated pattern on 49.2% of trials, and correctly recalled the associated scene on 82.1% of trials (both of which are well above chance performance [33.3%]). A paired-samples t-test revealed that memory for scenes was substantially greater than memory for patterns (p < 0.001, Cohen’s dz = 4.31). To attenuate the impact of differing memory performance for the two stimulus types in the subsequent analyses, two regressors were included in all models that served to suppress variance attributable to scene-specific and feature-specific memory.

### Neocortical alpha/beta power decreases during sequence perception predict enhanced memory performance

After establishing that alpha/beta power did indeed decrease from baseline following the presentation of the sequence stimuli (p_corr_ < 0.001, Cohen’s d_z_ = 1.22, cluster size = 28,260, mean t-statistic within cluster = -5.03; see figure 2a/d), we set out to test our first hypothesis: are memory-related decreases in alpha/beta power more prevalent during the perception of the sequence than during mnemonic binding?

**Figure 2.**
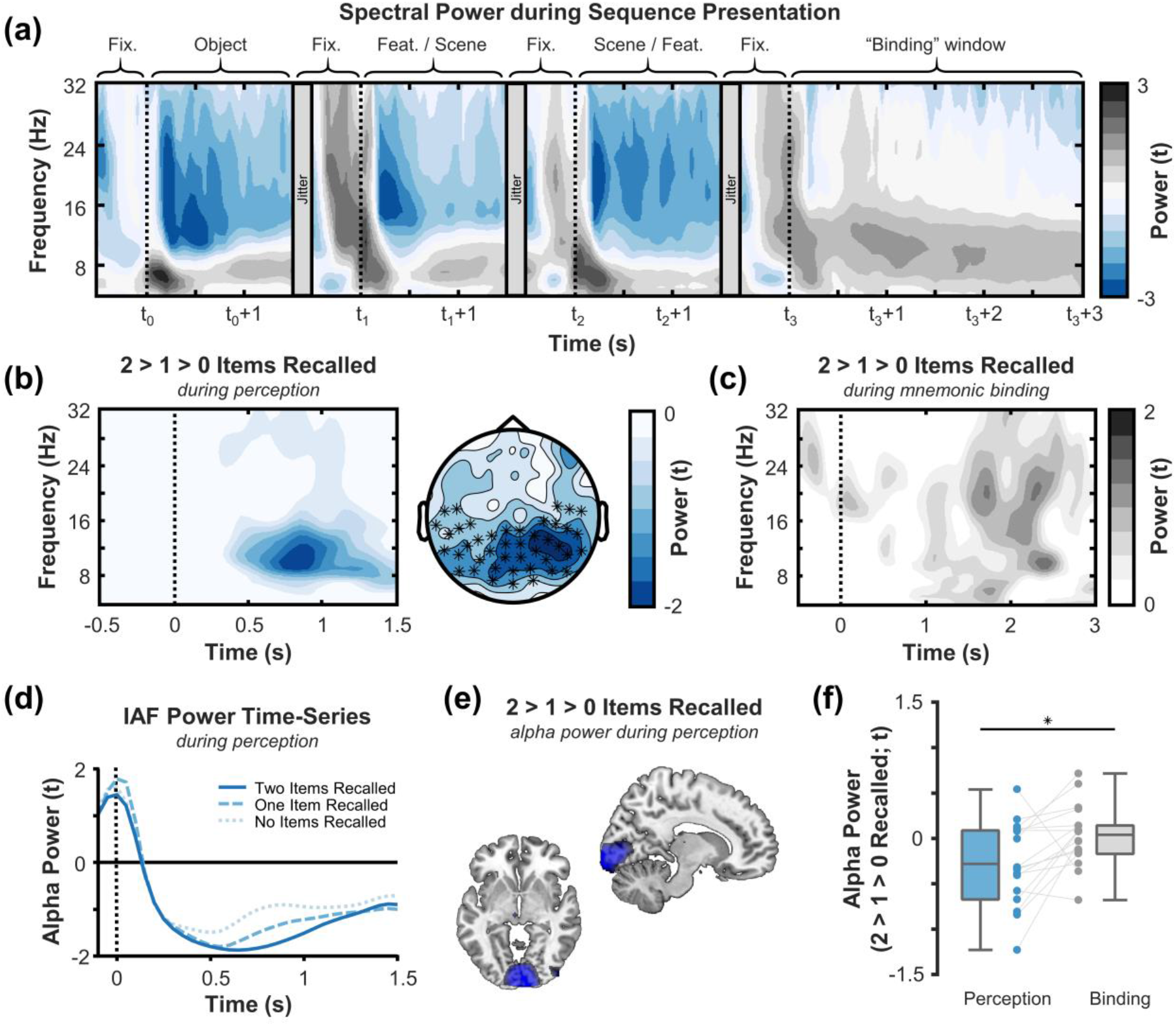
Neocortical alpha/beta power decreases during sequence perception scale with the number of items later recalled. **(a)** Time-frequency plot of alpha/beta power (averaged across all trials) during sequence presentation and subsequent binding. Alpha/beta power only decreases during sequence presentation. **(b)** Time-frequency plot (left) and topoplot (right) of the negative correlation between alpha/beta power during sequence perception and the number of items later recalled. The time-frequency plot uses the average of all channels included in the significant cluster (visualised by crosses in the topoplot to the right). The topoplot depicts values for time-frequency bins included in the significant cluster (i.e., 8-15Hz; 300-1300ms). **(c)** Time-frequency plot of the correlation between alpha/beta power during mnemonic binding and the number of items later recalled, plotted over the same channels as those visualised in panel b. No significant effect was observed. **(d)** Time-series plot of the power at the individual alpha frequency (IAF) of each participant for each memory condition. The more items later recalled, the greater the power decrease [pcorr = 0.034 at IAF]. **(e)** Source localisation of the effect in panel b. The memory-related alpha/beta power decreases during sequence perception peak in the occipital cortex. **(f)** Boxplot of memory-related decreases in alpha/beta power during sequence perception and mnemonic binding. Across participants, memory-related decreases in alpha/beta power were significantly greater during sequence perception than during mnemonic binding.

During sequence perception, cluster-based analysis revealed a significant effect where a decrease in alpha/beta power correlated with an increase in the number of items later recalled (p_corr_ = 0.032, Cohen’s d_z_ = 0.60, cluster size = 1013, mean t-statistic within cluster = -2.47; see figure 2a-f). This cluster extended over the posterior sensors, bilaterally, between 8 and 15Hz (see figure 2b). Source reconstruction confirmed this localisation, implicating bilateral early occipital regions (see figure 2e). Parsimonious results were found during retrieval (see supplementary figure 2), and when breaking the memory effects down by stimulus type (see supplementary figure 3).

No memory-related changes in theta power (2-7Hz; p_corr_ = 0.101), “slow” gamma power (40-60Hz; no cluster formed), or “fast” gamma power (60-100Hz; no cluster formed) were observed during the presentation of the sequence.

No decrease in alpha/beta power was observed when participants were asked to engage in mnemonic binding (p_corr_ = 0.413; see figure 2g). Similarly, no memory-related changes in theta power (2-7Hz; p_corr_ = 0.130), “slow” gamma power (40-60Hz; no cluster formed), or “fast” gamma power (60-100Hz; no cluster formed) were observed when participants were asked to engage in mnemonic binding.

A direct contrast of spectral power between sequence perception and mnemonic binding demonstrated that the inverse relationship between alpha power and subsequent memory performance was significantly more pronounced during perception (p_corr_ = 0.014, Cohen’s d_z_ = 0.60, cluster size = 794, cluster t-statistic = -2.49; see figure 2f). Together, these findings suggest that alpha/beta power decreases during sequence perception, but not during mnemonic binding, scale with the number of items that are later recalled.

### Hippocampal theta/gamma phase-amplitude coupling during mnemonic binding, but not information representation, predicts successful episodic memory formation

We then probed how hippocampal theta/gamma phase-amplitude coupling relates to episodic memory formation. During mnemonic binding, increases in hippocampal theta/gamma phase-amplitude coupling scaled with the number of items later recalled (t(16) = 2.24, p = 0.020, Cohen’s d_z_ = 0.54; see figure 3). No significant coupling was observed during perception (p > 0.5). A direct contrast in PAC between sequence perception and mnemonic binding revealed that memory-related increases in PAC are more pronounced during mnemonic binding (t(16) = 1.93, p = 0.040, Cohen’s d_z_ = 0.47). These results suggest that memory-related theta/gamma phase-amplitude coupling is most prominent during periods of mnemonic binding.

**Figure 3.**
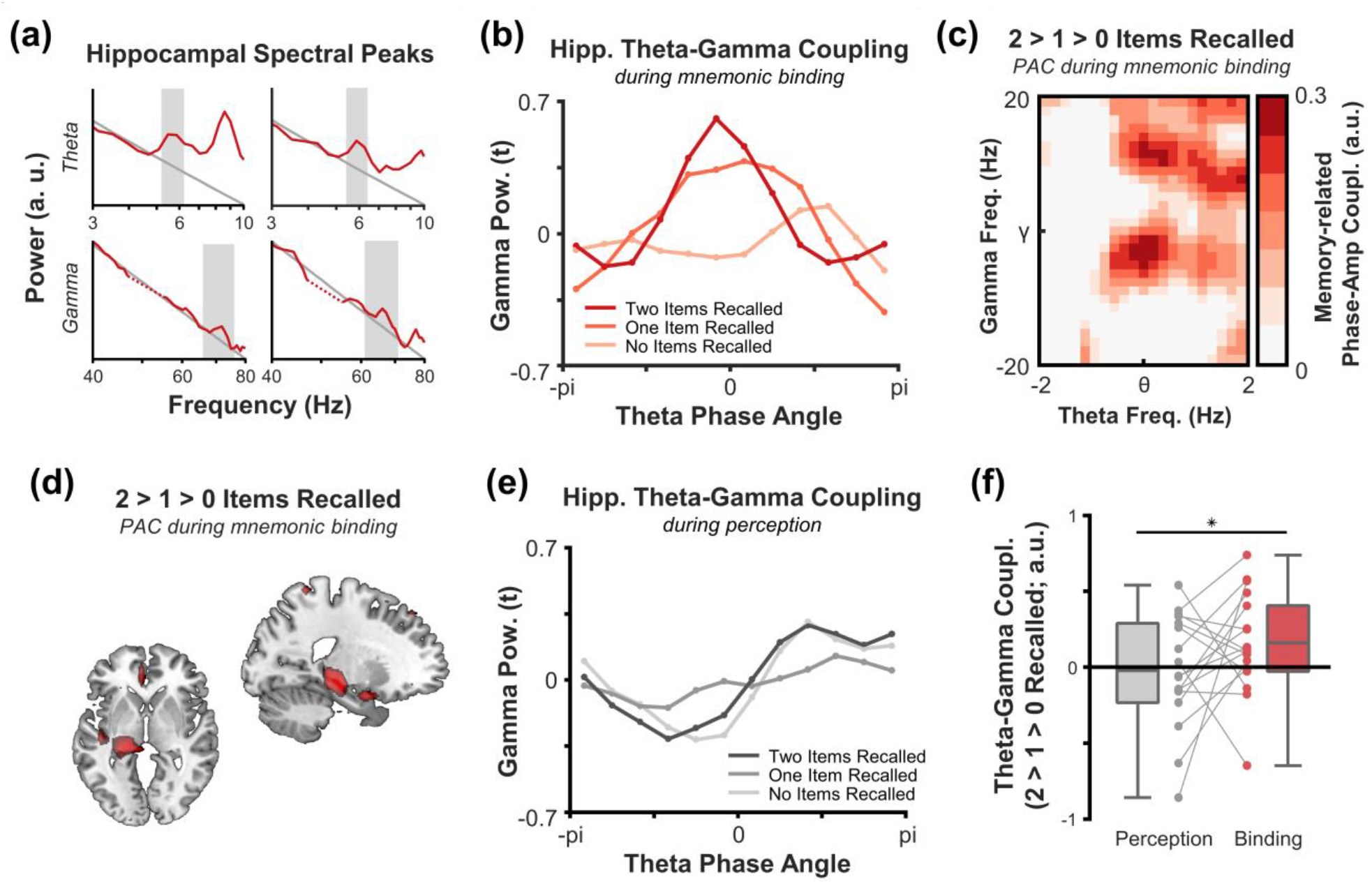
Increases in hippocampal theta-gamma coupling during mnemonic binding scale with the number of items later recalled. **(a)** Exemplar plots of peak theta and gamma frequencies for two participants (red line depicts hippocampal power; dotted red line depicts frequencies interpolated due to line noise; grey line depicts fitted 1/f component; grey area depicts identified peak). **(b)** Hippocampal gamma power as a function of hippocampal theta phase, for each memory condition, during the mnemonic binding window. When more items were later recalled, gamma power fluctuated in line with theta phase more noticeably. **(c)** Memory-related hippocampal theta-gamma coupling as a function of theta and gamma frequencies. Theta-gamma coupling appeared to peak at approximately the peak theta and gamma frequencies, supporting the idea that this coupling arises between two narrow-band oscillatory signals. **(d)** Theta-gamma coupling peaked in the hippocampus. To emphasise coupling patterns that were consistent across both hemispheres, the two hemispheres have been averaged together, and the averaged result visualised in the left hemisphere of the source plot. **(e)** Hippocampal gamma power as a function of hippocampal theta phase, for each memory condition, during sequence perception. No memory-related differences in coupling were observed. **(f)** Boxplot of memory-related increases in theta/gamma coupling during sequence perception and mnemonic binding. Across participants, memory-related increases in theta/gamma coupling were significantly greater during mnemonic binding than during sequence perception.

To ensure that the hippocampal effect observed during mnemonic binding was not a result of spatial smearing from some other region, we re-ran this analysis using four additional regions of interest: the frontal lobe, parietal lobe, temporal lobe (excluding the hippocampus), and the occipital lobe. None of these regions exhibited significant theta-gamma phase-amplitude coupling during mnemonic binding (frontal: p = 0.308, parietal: p = 0.250, temporal: p = 0.078, occipital: p = 0.169). Furthermore, a searchlight-based analysis revealed that hippocampal phase-amplitude coupling was substantially greater than other searchlight-based regions-of-interest that matched the size of the hippocampus (p = 0.024). Together, these results suggest that the memory-related enhancement in hippocampal theta/gamma phase-amplitude coupling is indeed originating from the source-reconstructed hippocampus, as opposed to “bleeding in” from neighbouring regions.

As can be seen in figure 3b, there is an apparent memory-related shift in the phase at which gamma couples to theta during menomnic binding. As the modulation index used above is insensitive to such shifts, we statistically appraised this effect using a circular-to-linear correlation. Alas, no consistent change was found (p = 0.435) suggesting that gamma activity does not precess along the phase of theta as a function of the number of items later recalled.

### A double dissociation between the timing of memory-related decreases in neocortical alpha/beta power decreases and memory-related increases in hippocampal theta/gamma phase-amplitude coupling

Lastly, we formalised the distinction between memory-related neocortical alpha/beta power decreases and hippocampal theta-gamma phase-amplitude coupling. To this end, we conducted a 2×2 repeated measures ANOVA with encoding stage (perception vs. binding) and metric (alpha/beta power decreases vs. theta-gamma coupling) as factors. This revealed a significant interaction where memory-related alpha/beta power decreases and memory-related theta-gamma coupling increases are dependent on the nature of the ongoing cognitive task [F(1,16) = 9.14, p = 0.008, partial eta squared = 0.36] (see figure 4). There was no main effect of cognitive task [F(1,16) = 0.99, p = 0.33, partial eta squared = 0.06], nor metric [F(1,16) = 0.20, p = 0.50, partial eta squared = 0.03]. These results, in conjunction with those reported in the sections above, suggest that a double dissociation exists between neocortical alpha/beta power decreases and hippocampal theta-gamma phase-amplitude coupling, with the former being most pronounced during periods of information processing and the latter being most pronounced during periods of mnemonic binding.

**Figure 4.**
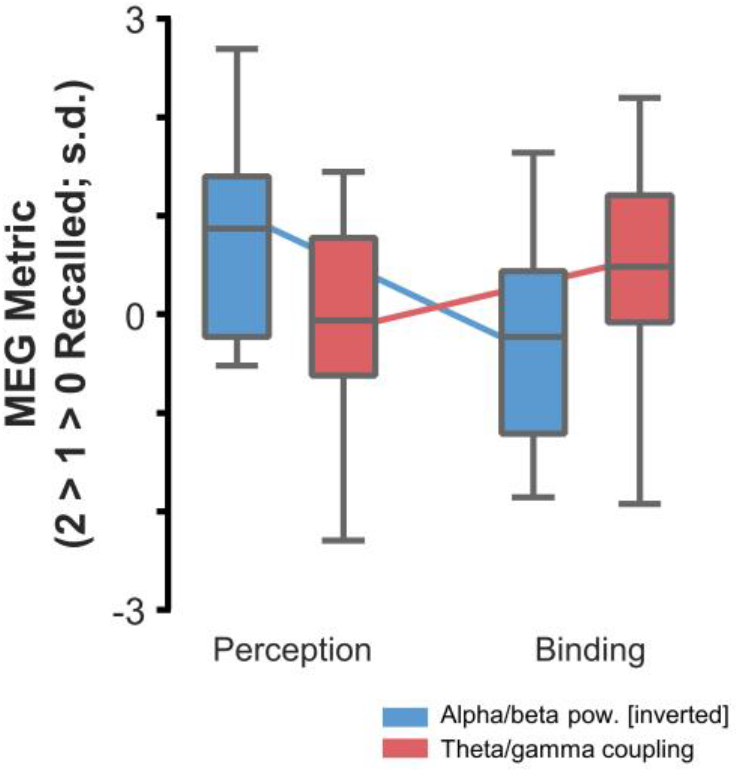
A temporal double dissociation between neocortical alpha/beta and hippocampal theta/gamma activity. Individual boxplots describe the effects of each task and each measure. The centre boxplot bar depicts the group mean, the box ends depict the 25^th^ and 75^th^ quartile and the tails depict the absolute minimum and maximum values within the sample. A significant interaction between the encoding stage and the MEG measure indicates that memory-related neocortical alpha/beta power decreases are most prominent during sequence perception, whereas memory-related increases in hippocampal theta/gamma coupling are most prominent during mnemonic binding.

## Discussion

Reductions in neocortical alpha/beta power and enhancements in hippocampal theta-gamma phase-amplitude coupling are thought to play dissociable roles in the formation of episodic memories (Hanslmayr et al., 2016), with the former supporting information representation and the latter supporting mnemonic binding. As such, one would expect that these neural phenomena are temporally dissociable, with alpha/beta power decreases arising first (supporting the processing of incoming information) and theta-gamma phase-amplitude coupling arising later (supporting mnemonic binding, which can only arise after the information has initially been processed in the cortex). Here, we found just that. Memory-related decreases in neocortical alpha/beta power principally arose during sequence perception, while memory-related increases in hippocampal theta/gamma phase-amplitude coupling principally arose during a time window in which participants could mnemonically bind the sequence together. This double dissociation suggests that alpha/beta power decreases and hippocampal theta/gamma phase-amplitude coupling arise at two distinct stages in the memory formation process.

The representation of information relating to a to-be-encoded memory is thought to be supported by neocortical alpha/beta power decreases (Hanslmayr et al., 2012). Information theory proposes that unpredictable states, such as desynchronised neural networks, carry more information than predictable states (Shannon & Weaver, 1949). In line with this hypothesis, we found that memory-related reductions in alpha/beta power (an index for neural desynchrony) only arose when participants were required to process information about a sequence, and not when the sequence was being bound together (i.e., when no further information was presented for processing). The restriction of alpha/beta power decreases to time points where information can be processed adds further support to the idea that alpha/beta power decreases correlate with the representation of information relevant to episodic memories (Griffiths, Mayhew, et al., 2019; Hanslmayr et al., 2012).

During mnemonic binding, hippocampal theta-gamma phase-amplitude coupling scaled with memory performance. Mechanistically speaking, these increases may reflect a heightened degree of long-term potentiation (LTP) within the hippocampus. By coupling gamma oscillations resonating at a frequency optimal for spike-timing dependent plasticity (STDP; Bi & Poo, 1998; Nyhus & Curran, 2010) to the phase of theta optimal for LTP (Hasselmo et al., 2002), the potential for building synaptic connections between hippocampal neurons is increased greatly. Based on such ideas, one could speculate that the memory-related theta-gamma coupling observed during the binding window reflects the transformation of the three discrete sequence stimuli into a singular cohesive episodic memory. Similarly, the absence of memory-related theta-gamma coupling during sequence perception may be attributable to the fact that the sequence is still unfolding, and, as such, a cohesive representation of all three stimuli cannot yet be formed. Notably, this is not to say that no theta-gamma coupling unfolds during sequence perception (indeed, several frameworks and empirical studies propose the opposite; Bahramisharif, Jensen, Jacobs, & Lisman, 2018; Griffiths & Fuentemilla, 2019; Heusser, Poeppel, Ezzyat, & Davachi, 2016; Lisman & Jensen, 2013), but rather, any theta-gamma coupling that does arise during that window is not predictive of later memory performance in this particular task. Perhaps instead, such coupling is predictive of item-context binding (e.g. Howard & Kahana, 2002), a measure not assessable here.

If these two phenomena are dissociable, does this mean that they act in complete independence of one another during encoding? Here, we would argue “no”. Mnemonic binding cannot occur if the relevant information has not been perceived, as there is no information to bind. Therefore, one could expect that the underlying neural correlates of mnemonic binding are contingent on the prior neural processing of relevant information. In line with such ideas, prior work has shown that the magnitude of hippocampal gamma synchronisation can be predicted by preceding neocortical alpha/beta power decreases (Griffiths, Parish, et al., 2019; see supplementary figure 4 for complementary findings within the data reported here). As such, one could speculate that neocortical desynchrony and hippocampal synchrony correlate with distinct cognitive processes (as evidenced above), but both neural phenomena (and the associated cognitive processes) must arise and interact to create an episodic memory.

We did not observe any memory-related fluctuations in theta power during the binding window. This is somewhat surprising; numerous previous studies have reported fluctuations in theta power correlating with later memory performance (for reviews, Herweg et al., 2019; Nyhus & Curran, 2010). Given that theories regarding theta and long-term potentiation (Bi & Poo, 1998; Hanslmayr et al., 2016; Nyhus & Curran, 2010) emphasise the importance of phase for LTP, rather than power, one could speculate that theta power has less to do with enhanced mnemonic binding, and as such, should not substantially correlate with successful memory formation. Similarly, we did not observe fluctuations in gamma power during mnemonic binding despite numerous studies demonstrating this previously (Burke et al., 2013; Griffiths, Parish, et al., 2019; Long & Kahana, 2015; Osipova et al., 2006). However, this can be explained by the fact that theta-gamma coupling was observed during this same window. If memory-related increases in gamma power are restricted to certain phases of theta, and theta is not stimulus-locked across trials, then across-trial averages of gamma power will sum to zero. As such, the absence of a ‘pure’ gamma effect here is not surprising.

In sum, these results demonstrate that decreases in neocortical alpha/beta power and increases in hippocampal theta/gamma phase-amplitude coupling are temporally dissociable in episodic memory formation (Hanslmayr et al., 2016).

## Supporting information

Response to Reviews

## Footnotes

* While it is impossible to conclude with absolute certainty that perception and mnemonic binding are completely separable in any memory task, here we can conclude that there is a substantial shift in the ratio of the two processes. Stimulus perception will only be taking place while there is stimulus to perceive (i.e., during the presentation of the sequence), while mnemonic binding will be most prevalent when all sequence items have been presented and processed (i.e., after the final stimulus has been processed by the cortex). While this leaves room for some binding to occur towards the very end of the presentation of the last stimulus, this would be minimal in comparison to what follows during the “binding window” (see figure 1a, outlined in red). Direct contrasts of the MEG signals between sequence perception and mnemonic binding will empirically test this hypothesised shift in ratio.

† Though the notion of localising deep regions such as the hippocampus was once controversial, an ever-growing number of studies have suggested that it is achievable. Ruzich and colleagues (2019) uncovered 29 studies that used gradiometers alone to localise hippocampal signals, while Dalal and colleagues (2013) demonstrated that MEG signals directly correlate with simultaneously recorded intracranial hippocampal recordings. As such, the theoretical notion that the hippocampus cannot be measured using MEG has been refuted by numerous empirical demonstrations to the contrary.

