## Supplementary Materials for "Disentangling neocortical alpha/beta and hippocampal theta/gamma oscillations in human episodic memory formation"

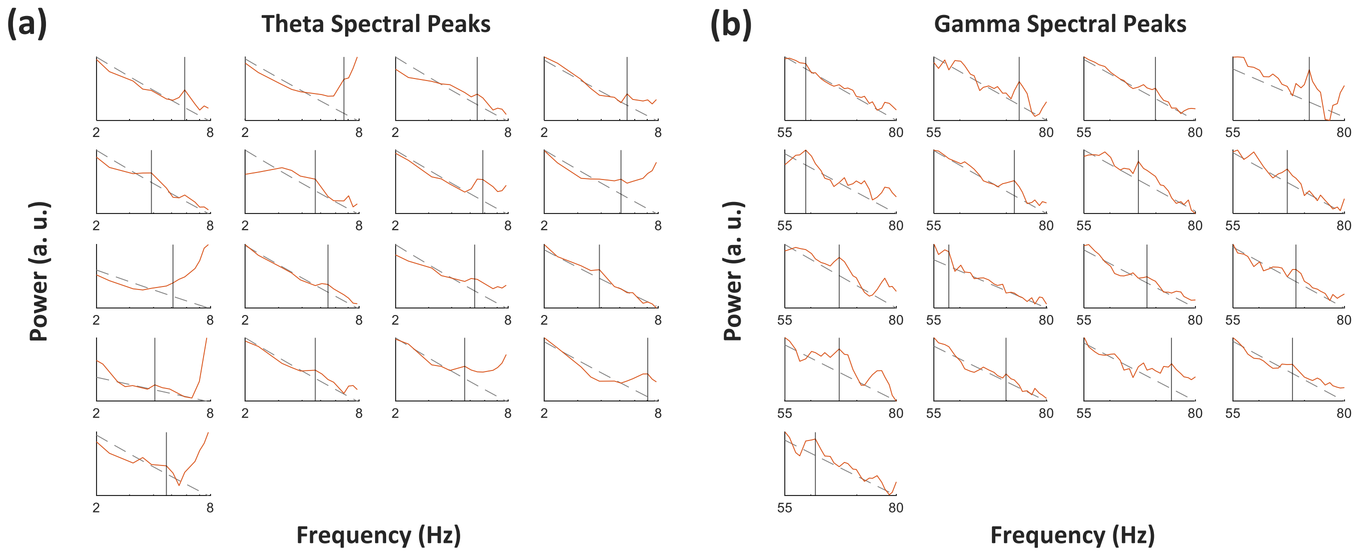


**Supplementary Figure 1.** Spectral peaks in the hippocampus. **(a)** Theta spectral peaks (red line: spectral power; dotted grey line: linear fit; black line: detected peak). Note that in some instances (sub-02, sub-08 and sub-09), the peak was embedded in the cortical alpha peak. Therefore, the identified peaks were cross-checked by taking the first derivative of the plotted signal and confirming that the derivative peak overlapped with the originally detected peak. **(b)** Gamma spectral peaks (red line: spectral power; dotted grey line: linear fit; black line: detected peak)

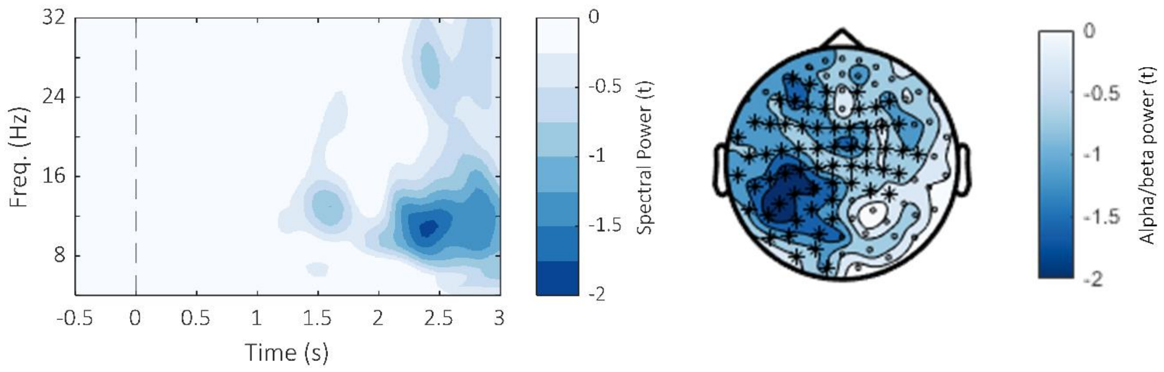


**Supplementary Figure 2.** In the main text, it is proposed that alpha/beta power decreases scale with the amount of information present in the cortex – a hypothesis that can also be generalised to the retrieval data (i.e., decreases in alpha/beta power during memory retrieval correlate with the number of items recalled). Indeed, when testing this hypothesis, we found that alpha/beta power decreases during retrieval correlated with the number of items recalled ($p_{corr}$ = 0.027, Cohen's $d_z$ = 0.59, cluster size = 1311, mean t-statistic within cluster = -2.42). This supports our current hypothesis and conceptually replicates findings from similar experiments that have focused alpha/beta power during memory retrieval (Karlsson et al., 2020; Martín-Buro et al., 2020).

No memory-related change in hippocampal theta-gamma coupling was observed during memory retrieval (p = 0.161).

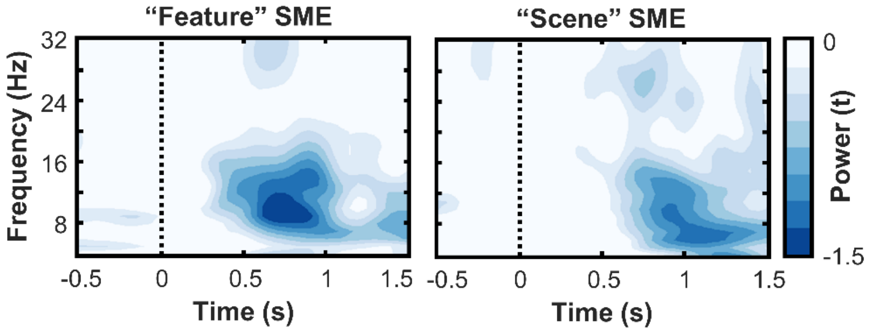


**Supplementary Figure 3.** Stimulus-specific memory effects. During the presentation of both feature (left) and scene (right) stimuli, the magnitude of the alpha/beta power decrease over occipital regions (plots use cluster reported in the main text) predicted later recall success [feature: $p_{clus}$ = 0.003; scene: $p_{clus}$ = 0.061]. It’s worth noting that memory performance was substantially better for scene stimuli than feature stimuli, leading to a “hit vs. miss” trial imbalance in the former that limits its statistical power.

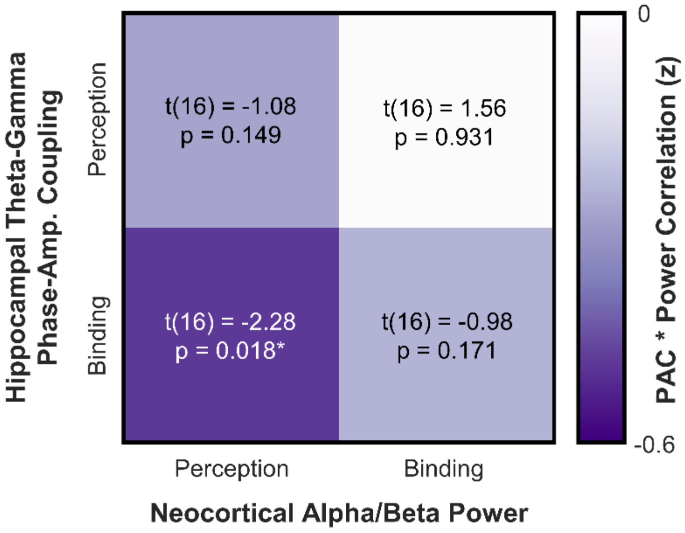


**Supplementary Figure 4.** Neocortical alpha/beta power decreases during perception correlate with increases in hippocampal theta-gamma coupling during mnemonic binding on a trial-by-trial level. No effect was observed when correlating these metrics at other epoch combinations (e.g., correlating alpha/beta power during binding with theta/gamma coupling during binding).

For each participant, a linear model [consisting of (1) a constant, (2) neocortical alpha/beta power during perception (in the cluster described in the main text), and (3) number of items recalled] was used to predict hippocampal theta-gamma coupling during mnemonic binding. The second co-efficient (i.e., alpha/beta power) for each participant was standardised, and then pooled for a one-tailed t-test (assuming a negative relationship between neocortical alpha/beta power and hippocampal theta-gamma coupling). Indeed, we found that the more that alpha/beta power decreased during sequence perception, the more hippocampal theta-gamma coupling increased during the following binding window [$t(16) = -2.28$, $p = 0.018$]. Note that this cannot be attributed to a phantom correlation induced by both MEG metrics correlating with the number of items recalled, as this source of variance was accounted for in the participant-specific linear models.

No similar effect was observed for any other combination of power*PAC correlations.

These results match Griffiths and colleagues' (2019) findings that neocortical alpha/beta power decreases precede hippocampal gamma power increases during episodic memory formation.
