## Supplementary material for "Disentangling neocortical alpha/beta and hippocampal theta/gamma oscillations in human episodic memory formation": Response to Reviews

*The following reviews were provided by eLife and are publicly available on bioRxiv.*

<u>General Response to Reviewers</u>

We would like to thank the reviewers for taking the time review our manuscript. The comments below have been thought-provoking and will inspire several new analyses that we hope address concerns. In particular, we will carefully reappraisal the framing of the results, shifting away from a false dichotomy of "this is perception" and "this is binding", and towards more restraint terminology that discusses the shift in balance between perception and binding. Moreover, we will expand our analysis of theta-gamma phase-amplitude coupling beyond the hippocampus and to the whole brain.

We answered each comment in turn, first by providing a general response to the comment and then by providing an outline of the explicit action we will take to address this issue.

<u>Reviewer #1</u>

**This MEG study by Griffiths and colleagues used a sequence learning paradigm which separates information encoding and binding in time to investigate the role of two neural indexes - neocortical alpha/beta desynchronization and hippocampal theta/gamma oscillation - in human episodic memory formation. They employed a linear regression approach to examine the behavioral correlates of the two neural indexes in the two phases, respectively and demonstrated an interesting dissociation, i.e., decreased alpha/beta power only during the "sequence perception" epoch and increased hippocampal theta/gamma coupling only during the "mnemonic binding" phase. Based on the results, they propose that the two neural mechanisms separately mediate two processes - information representation and mnemonic binding. Overall, this is an interesting study using a state-of-art approach to address an important question. Meanwhile, I have several major concerns that need more analysis and clarifications.**

<u>Major comments:</u>

**(1) The lack of theta-gamma coupling during stimulus encoding period is possibly due to the presentation of figure stimulus, which would elicit strong sensory responses that mask the hippocampus activity. How could the author exclude the possibility? In other words, the dissociated results might derive from different sensory inputs during the two phases.**

> <u>Response:</u> The reviewer raises a good point; However, we feel this is already addressed by our use of memory-related contrasts. The masking of an effect that arises due to stimulus presentation would be consistent across all memory conditions, and therefore subtracted out in any contrast between these conditions. The analyses in our original submission use this approach to avoid such a confound. Furthermore, previous studies (e.g., Heusser et al., 2016, Nat. Neuro.) have demonstrated that hippocampal theta-gamma coupling can arise during stimulus presentation, suggesting strong sensory responses do not, generally speaking, mask measures of theta-gamma coupling.

**(2) About the hippocampal theta/gamma phase-power coupling analysis. I understand that this hypothesis derives from previous research (e.g., Heusser et al., 2018) as well as the group itself (Griffiths et al., PNAS, 2019). Meanwhile, MEG recording, especially the gradiometer, is known to be relatively insensitive to deep source. Therefore, the authors should provide more direct evidence to support this approach. For instance, the theta/gamma analysis relies on the presence of theta-band and gamma-band peak in each subject. Although the authors have provided two**

**representative examples (Figure 3A), it remain unknown how stable the theta-band and gamma-band peak exist in individual subject.**

Response: Regarding concerns about the MEG gradiometers being relatively insensitive to deep sources, we feel it is worth noting that a recent review (Ruzich et al., 2019, Human Brain Mapping) identified 29 studies that had reported successful hippocampal measurements when only using gradiometers, suggesting our use of gradiometers is not unprecedented nor unjustified. Furthermore, in their recommendations for optimising hippocampal recordings with MEG, the old wisdom of using magnetometers rather than gradiometers is conspicuous in its absence in the review – perhaps because while magnetometers have a greater theoretical potential to detect deep signal, they also have greater theoretical potential to pick up noise, so the signal-to-noise ratio (which, arguably, is key here) for deep sources may not differ so much between gradiometers and magnetometers.

Action: We have added a supplementary plot that visualises the theta/gamma band peaks for each participant (see supplementary figure 1). We have also discussed the feasibility of using MEG to measure deep sources in the methods (see footnote on page 7).

**(3) Related to the above comment, the theta-gamma coupling is a brain-wide phenomenon including both cortical and subcortical areas and not limited to just hippocampus. Although the authors have performed a control analysis to assess the behavioral correlates of the coupling in other regions, the division of brain region is too coarse and I am not convinced that this is a fair comparison, since they differ from hippocampus at least in terms of area size in the source space. The authors could consider plotting the power-phase coupling distribution in the source space and then assessing their behavioral correlates, rather than just showing results from hippocampus. This result would be important to confirm the uniqueness of hippocampus in this binding process.**

Response: We concur that the plots currently do not demonstrate the specificity of the hippocampus, and whole brain images would better demonstrate the effect.

Action: As suggested by the reviewer, we have plotted theta-gamma coupling across the brain (see figure 3d). We have also compared hippocampal PAC to searchlights that match the size of the hippocampal ROI, and found that hippocampal PAC is significantly greater than what is observed in these searchlights (see lines 276-287).

**(4) About behavioral correlates. The current behavioral index confounds encoding and binding processes. Is there any way to seperate the encoding and binding performance from the overall behavioral measurements? it would be more convincing for me to find the two neural indexes at two phases predict the two behavioral indexes, respectively.**

Response: This is a really interesting idea, but one which perhaps requires a different experiment paradigm. For associative memory, we would argue that binding is an essential step for the successful encoding of a memory, so it would quite possibly be impossible to separate the two processes in the paradigm used here. That said, a different paradigm that compared associative memory to, say, item memory, may be able to answer such a question.

**(5) The author's previous works have elegantly shown the two neural indexes during fMRI and intracranial recording in episodic memory. The current work, although provideing interesting view about their possible dissociated functions, only focuses on the memory formation period (information encoding and binding). Given previous works showing interesting relationship between encoding and retrieval (Griffith et al., PNAS, 2019), I would recommend the authors to also**

**analyze the retrieval period and see whether the two indexes show consistent dissociated function as well.**

Response: Yes, we completely agree. We had included this in a previous draft of the manuscript, and found a consistent dissociation here, where alpha/beta power decreases accompanied retrieval (perhaps linked to the representation of retrieved information) and theta-gamma coupling did not (perhaps due to the absence of a need to bind stimuli together to complete the retrieval task).

Action: We have added these results to the supplementary materials to maintain concision in the manuscript (see supplementary figure 2).

**Minor:**

**(1) It is not clear to me how the channels showing behavioral correlates in alpha/beta power are selected (Figure 2B). Are the statistical tests corrected across channels?**

Response: The channels highlighted in figure 2b are those which belong to the significant cluster.

Action: We have clarified the channel selection in the figure legend.

**(2) The Figure 2a legend reads "...depicting fluctuation in power as a function of number of items recalled...". How does the figure illustrate the behavioral correlates, and isn't it just plotting the time-frequency power? The same question for Figure 2b and Figure 3c.**

Response: The metric plotted is the standardised beta coefficient which describes how spectral power can be predicted by the number of items recalled, rather than raw spectral power. This is a way to plot a contrast in spectral power (e.g., A > B) when there are more than two conditions (as is the case here: two items recalled > one item recalled > no items recalled).

Action: We have clarified the measure plotted in the TFR in the figure legend and re-labelled the plot to avoid confusion.

**(3) The authors performed a regression analysis on the neural indexes according to recalling performance (number of item recalled). If the conclusion is right, we would expect a linear decrease in alpha/beta power during the first phase and a gradual increase in theta-gamma coupling during the 2nd phase, as the number of recalled item increases. This could be explicitedly plotted.**

Response: Yes, we would expect a monotonic decrease in alpha/beta power during the first phase of the task, and a monotonic increase in theta/gamma power during the second phase.

Action: We have plotted alpha/beta power and theta/gamma coupling for the three memory conditions separately (see figures 2d and 3b).

**Reviewer #2:**

**In this manuscript, the authors examine the neural correlates of perception and memory in the human brain. One issue that has plagued the field of memory is whether the neural processes that underlie perception can be dissociated from those that underlie memory formation. Here the authors directly test this question by introducing a behavioral paradigm designed to dissociate perception from mnemonic binding. In brief, while recording MEG data, they present subjects with**

**a sequence of visual stimuli. Following the sequence, the subjects are instructed to bind the three stimuli together into a cohesive memory, and then are tested on their memory for which pattern was associated with an object, and which scene. The authors investigate changes in alpha/beta power and theta/gamma phase amplitude coupling during two separate epochs - perceptual processing and mnemonic binding. Overall, this is a well written and clear manuscript, with a clear hypothesis to be tested. Using MEG data enables the authors to draw conclusions about the neurophysiological changes underlying both perception and memory, and establishing this dissociation would be an important contribution to the field. I think the conclusions are justified, but there are several issues that should be addressed to improve the strength and clarity of the work.**

**The fundamental premise of the task design is that subjects view a sequence of stimuli, and then separately at a later time actively try to bind those visual stimuli together as a memory. However, it is entirely possible, and even likely, that memories are being formed and even bound together as the subjects are still viewing the sequences of objects. How would the authors account for this possibility? One possible way would be if there were a control task where subjects were just asked to view items and not remember them.**

> <u>Response:</u> Indeed, it is impossible to ascertain that no binding is occurring during sequence presentation, and the terminology used in the original submission is ill-fitting as a result. However, we would argue that there is a shift in the ratio between perception and binding across the encoding task, with greater perceptual processes arising during the presentation of the sequence relative to the "associate" cue (as this is when the items are presented), and greater associative processes arising during the "associate" cue (as this is when all items are available for binding). To suggest that the two processes can be completely separated would be erroneous, but we feel it is also difficult to argue that there is no shift in balance between the two processes over the course of the encoding task. Importantly, linking a shift in balance between the two processes (binding/perception) with neurophysiological correlates (alpha-beta/theta-gamma) is sufficient to support our main conclusion.
>
> <u>Action:</u> We have carefully rephrased the manuscript in such a way that it no longer implies that there is a perfect separation of perception and binding, but rather a shift in the balance between the two processes. We have also explicitly addressed this in a footnote to the final paragraph of the introduction (see page 1), where we state that it is impossible to conclude that the processes are perfectly separate, but nonetheless show a substantial shift in the ratio of the processes.
>
> <u>Note on a "control" task:</u> In our view, the control task proposed by the reviewer is captured by the "forgotten" condition – participants view the items, but do not subsequently remember them.

**Another possibility would be to examine the trials that the participants failed to remember correctly. Presumably, one would still see the same decreases in alpha power. Yet it seems from the data, and the correlations, that during those trials that were not remembered properly, alpha power change very little. Of course, it is unclear in these trials if failed memory is due to failed perception, but one concern would that this would imply that decreases in alpha power are relevant for memory too. It would be helpful to see how changes in alpha power break down as a function of the number of actual items remembered. It would also be helpful to know how strong these correlations actually are.**

<u>Note:</u> We are a little unsure of what the reviewer is suggesting here, as we feel that most of these analyses were included in the main text. The response below re-cap of the results and how they link to our interpretation of the reviewer's comment, but if we have misunderstood the point, we would be willing to re-address it.

<u>Response:</u> In the original submission, we had focused solely on the memory-related change in alpha/beta power (that is: the contrast "2 items recalled" > "1 item recalled" > "no items recalled"). Therefore, the inferential statistics allow us to conclude that a relative decrease in alpha/beta power correlates with an increase in number of items recalled. What the analyses in the original submission do not show is that alpha/beta power changes from baseline (that is, are all items perceived [i.e., as indexed by a power decrease], or just the remembered items?). This is something, however, we have now included.

<u>Action:</u> We have reported the stimulus-induced change in alpha/beta power during sequence perception (see page 7-8, lines 282-284; see figure 2a and 2d).

**A related issue is with respect to hippocampal PAC. The authors investigate this during the mnemonic binding period. Yet they also raise the possibility in discussion that this could also be happening during perception, which goes back to the point above. Did they analyze these data during perception, and are there changes with perception that correlate with memory? This would suggest that binding is actually occurring during this sequence of visual stimuli.**

<u>Response:</u> We did indeed analyse the data during perception in the original submission (see lines 127-128; figure 3d) and found no evidence to suggest that memory-related PAC varied during perception. In an additional analysis, we also examined with PAC varied as the sequence progressed (that is, does PAC change from the first item to the second, and from the second to the third?), but found no evidence to suggest it does. Together, these results would suggest that putative binding mechanisms are not dominating the sequence perception phase of encoding.

**The authors perform a whole brain analysis examining the correlation between alpha power and memory to identify cluster corrected regions of significant. However, the PAC analysis focuses only on the hippocampus, raising the question of whether these results can account for the possible comparisons one could make in the whole brain. They do look at four other brain regions for PAC, which it would be helpful to account for. In addition, are there other measures of mnemonic binding that are significant? For example, theta power, or even gamma power?**

<u>Response:</u> We had focused our PAC analyses on the hippocampus because of our a priori hypotheses, but appreciate that only showing data from the hippocampus would obscure the whole picture. Our analyses did not uncover convincing evidence for changes in theta or gamma power, but we will report these in the main text and in the discussion

<u>Action:</u> We have presented the PAC results across the whole brain (see figure 3d), have reported analyses of theta and gamma power (see page 9, lines 303-306), and have discussed the absence of theta/gamma power effects in the discussion (see page 12, lines 397-409).

**The authors note in the discussion that the magnitude of hippocampal gamma synchrony has been shown to be related to the decreases in alpha power. Is this also true in their data?**

<u>Action:</u> We have added supplementary analysis (see supplementary figure 4) probing the correlation between hippocampus theta/gamma activity and neocortical alpha/beta power. In line with our earlier findings, we find that alpha/beta power decreases precede hippocampal theta-gamma increases.

**Reviewer #3:**

**The authors report results of an MEG analysis deploying a cognitive paradigm in which participants engage in a source memory task characterized by the appearance of three images in succession and are then tested via a cue (the first of the three images) followed by a choice of responses for a two dimensional pattern and then a choice (out of three images) of a photographic scene.**

**The principal finding is that (via MEG sensor level data) there is a widespread 8-15 Hz power decrease that is correlated with the number of recalled items (from 0 to 2) on a given trial. In the hippocampus (via MEG source reconstruction), the magnitude of phase amplitude coupling observed as participants are told to associate the items is correlated with memory performance. The 8-15 Hz power decrease/memory correlation (as estimated by beta coefficients in a model described in Figure 1) is larger (across individuals) during moments when subjects are viewing the stimulus items as opposed to during the "associate" period. The novelty in the result is related to the experimental task that attempts to dissociate memory-related effects related to perception from those related to binding which putatively occurs when subjects are given the "associate" instruction.**

**My main conceptual concern is related to the design of the experimental task. I am not sure that the perception/binding framing is appropriate, since there is not reason to think that subjects are not associating/binding items during the periods when the items are being shown on the screen. I suppose this may partly explain the lack of a significant difference in PAC/memory beta coefficients observed in the hippocampus when contrasting these two epochs (Figure 4). But the corollary is that the alpha power-related beta coefficients are observed while binding is likely also occurring within the paradigm (esp since each image is shown for 1.5 seconds it would seem). Is the alpha power effect seen in the hippocampus? The plots in 3a suggest there is an oscillation present in the relevant frequency range, and the time course of alpha power differences seen in Figure 2 suggests that they occur relatively late after onset of the images, which may fit better with some contribution for this pattern to the forming of associations rather than perception.**

Response to comments on task: We agree that the task does not unequivocally separate the two cognitive tasks, and any statement to suggest that it does is erroneous. That said, we would argue that, on a balance of probability, there is likely to be more information processing going on during sequence perception relative to when the associate cue is presented. This is because the participant is still being exposed to rich stimuli during sequence presentation, while only being presented with a simple cue during the association phase. Similarly, there is likely to be more binding during the associate cue than during sequence presentation. This is because participants have greater cognitive resources available for binding during the associate cue relative to during sequence perception. Now, neither of these reasons are sufficient to argue that “association” does not occur during sequence perception. However, we feel that these reasons are sufficient to suggest we expect to see a shift in the balance of “association” between the sequence perception and the binding window, where “association” is more easily executed during the binding window. Indeed, we feel it would be difficult to argue that there is no shift in the balance between these processes at any point. Importantly, linking such a shift in balance between the two processes (binding/perception) with neurophysiological correlates (alpha-beta/theta-gamma) is sufficient for our main conclusion. As such, we feel a careful rephrasing can address these concerns, where portions of the text referring to a separation of perception and binding are rephrased as a “shift in the balance in perception and binding” – the latter phrasing allows for the possibility that there is some small mixing of the two tasks.

<u>Action to comments on task:</u> We have carefully rephrased the manuscript such that the text does not suggest that perception and binding are perfectly separated (we have also added a footnote on page 3 that directly addresses this concern). Now, it simply suggests that the balance between the two processes shift during the encoding task.

<u>Response to comments on hippocampal alpha:</u> We agree that there appears to be an alpha peak in the hippocampus, but as this plot is across all trials, and this alpha oscillation does not correlate with memory (see page 9, lines 294-297).

**I understand that the paradigm was constructed in an attempt to temporally dissociate memory effects attributable to perception versus those attributable to binding. But given the temporal resolution available using EEG, I would imagine that the authors could differentiate an earlier perception-related effect from a later PAC binding effect in the time series if the associated images were presented in conjunction. Is it correct to frame the alpha results as related to "perception?" The beta coefficients used for analysis reflect a "memory related effect observed when visual stimuli are present on the screen," but not necessarily improved memory predicated on more accurate perception to my interpretation. I would think that a perception/binding distinction requires operationalizing perception as activity that doesn't vary with later associative memory success, and binding as activity that does. The notion of perception used by the authors here seems slightly different. The authors can perhaps comment on this concern.**

<u>Response:</u> This is a very interesting point. A hallmark of visual perception is a reduction in alpha/beta power (e.g., Pfurtscheller et al., 1994, Int. J. Psychophysiology), regardless of whether it is remembered or not. As such, we would expect alpha/beta power to decrease following stimulus onset even if a memory is not formed. We can directly test this by examining the stimulus-evoked power decrease in all conditions, with the expectation that alpha/beta power drops from baseline in all conditions.

<u>Action:</u> We have contrasted pre-stimulus and post-stimulus power and, indeed, found that alpha/beta power decreases accompany visual perception regardless of whether the memory was successfully encoded (see page 7, lines 282-284; see figure 2a and 2d).

**The authors report PAC results for other regions on page 6, but claiming that PAC is a hippocampal-specific effect would require showing that the PAC-related beta coefficients are significantly greater than the other regions, rather than simply the absence of a significant effect in these regions. The authors should also clarify if they combined locally measured PAC over several ROIs into an average for these other regions? It seems unlikely to detect PAC if a single theta/gamma time series were extracted over such a large area of cortex.**

<u>Response:</u> We agree with the principle that the PAC results should be probed further, though would argue against the use of inter-region contrasts here as they will not provide evidence that PAC is specific to a single region. Take, for example, an effect where there is a significant memory-related increase in PAC in region A, but there is a significantly larger memory-related increase in region B. In a direct contrast, PAC in B will be significantly greater than A, but clearly PAC is not specific to B. Therefore, an inter-region contrast is not a means to irrefutably demonstrate regional specificity. While there has been a call for direct comparisons between experimental contrasts (see Nieuwenhuis et al., 2011), this is specifically for cases where individuals wish to make the claim that “A is significantly greater than B”, which was a claim that we never made. Rather, we asked whether there is a memory-related difference in PAC within the hippocampus, and then followed this up by confirming that this effect was not a

"bleed-in" from PAC in another neighbouring region (i.e. the cortical ROI analyses; where the absence of a significant difference would suggest that memory-related hippocampal PAC is not attributable to memory-related PAC in another region). We will, however, better visualise the PAC results to further rule out the risk of a "bleed-in" effect (see response to Reviewer 1, point 3).

Action: We have clarified the rationale for these ROI analyses, and have visualised PAC across the cortex. We have also run a searchlight-based analysis that demonstrates the hippocampal PAC is significantly greater than what is observed in placed in other regions of the brain searchlights (see page 7, lines 276-287).

**The interaction effect reported at the end of the results (ANOVA model) is interpreted such that the cortical alpha effect is stronger when the visual items are presented, while the hippocampal PAC effect is stronger when no items appear on the screen, but these recordings are made in different regions (hippocampus versus the entire cortex). If my understanding is correct, a result in line with the model the authors suggest (cortical alpha power decrease/hippocampal PAC) would show a region (hipp v cortex) x task (images on screen vs "associate" command) x metric (PAC vs alpha) interaction. Can the authors clarify if the cortical data entered into this model includes only those regions that showed a significant effect initially, or just all the sensors? The former would seem to introduce bias.**

Response: Yes, for computational simplicity, we had selected the regions based on what was previously significant. By and large, the goal of the last section was not to provide any profound new insight into the data, but rather place the cortical power and hippocampal PAC results side by side (visually, in the case of figure 4) so one can appreciate the juxtaposition that has been uncovered in the previous sections. The statistical results we reported were more of a pre-emptive strike against those who demand statistics with their figures, and have not been used to further the conclusions of the paper beyond what the earlier analyses could already do by themselves. As such, any bias in the analysis is without consequence for the conclusions.

**Similarly, the different visual classes are always presented in the same order, which may give rise to the strong disparity in recall fraction between the pattern and scene images. I understand the linear model incorporates predictor variables for scene/pattern recall, but given that scene recall is driving a significant amount of the overall recall number observed as the main variable of interest, I would wonder if the alpha/beta power effects are related to the relative complexity of the scene images as compared to the patterns. Given the analysis schematic the authors report, I assume the authors have analyzed whether the same effects occur when contrasting scene versus no recollection and pattern vs no recollection. If the same effects are observed regardless of type of image (when compared with no recollection) this may help address this concern.**

Action: We have included supplementary analyses (see supplementary figure 3) which demonstrate that alpha/beta power decreases predict memory formation regardless of stimulus type.

Additional note: The scene and pattern stimuli were not always presented in the same order, but rather counterbalanced across blocks to avoid order effects.

**My second conceptual question is related the MEG data. It appears to me that the authors use MEG sensor-level data for the alpha-related effect in the cortex (Figure 2), but MEG beamformer reconstructed data (localized to the hippocampus) for the PAC effect. Is there a reason the authors did not use MEG data localized to specific cortical regions rather than sensor data? This may reflect**

**confusion on my part, but I don't understand why they would use qualitatively different types of data for these two aspects of the analysis that are then combined (in the ANOVA, for example).**

Response to questions on source-reconstructed alpha power: We had not included source-reconstructed analysis of the alpha power effect here because, in an earlier draft, extensive analysis (e.g., the reporting of both sensor-level and source-reconstructed alpha power effects) drew criticism from reviewers for a lack of conciseness. That said, it does make more sense to include such analyses.

Action: We have included source-reconstructed alpha-band effects (see figure 2e).

**The authors should also engage with concerns regarding the validity of localizing MEG signals (especially for an analysis such as PAC) to deep mesial temporal structures such as the hippocampus. I understand that MEG systems with greater than 300 sensors are more reliable for this purpose, but I think a number of readers would still have doubts about MTL localization of signal. Also, my understanding is that such deep source localization requires around 100 trials per class, which I think fits with what the subjects completed, but the authors may include references related to this issue.**

Response: In recent years, there has been a growing list of studies that have reported successful localisation of hippocampal signals using MEG (for review of 37 of these studies, see Ruzich et al., 2019, Human Brain Mapping). Generally speaking, our experimental paradigm and analysis pipeline show large overlap with these previous successes (e.g., use of beamformers, gradiometers, co-registered MRI-to-MEG head position), meaning our results are not completely out of line with what could be expected.

Action: We have explicitly addressed the historic difficulties of localising hippocampal MEG signals, and highlight how our approach fits with a growing consensus on how to successfully localise such signals (e.g., Ruzich et al., 2019, Human Brain Mapping) [see footnote on page 7].

**I think the signal processing steps are overall quite reasonable. I would ask the authors to clarify if they limited their analysis of cortical alpha/beta oscillations to those in which a peak exceeded the 1/f background, as they report for the PAC analysis on page 5. Also, it would be helpful to show that the magnitude of the MI values in the hippocampus exceed those observed by chance (using a shuffle procedure) in addition to showing that there is a memory-related association reflected in the beta coefficients.**

Response: We had not limited the analysis to peak alpha/beta oscillations in the original submission, but it is an interesting question.

Action: We have restricted alpha/beta power analysis to the peak oscillation, and the results remain the same (see figure 2d). We have added supplementary analysis contrasting measures of hippocampal PAC to a shuffled baseline.
